# A recombinant hybrid provides insights into gene regulation, pathogenesis and tumorigenesis of phytopathogenic smut fungi

**DOI:** 10.1101/2025.01.28.635268

**Authors:** Janina Werner, Weiliang Zuo, Tom Winkler, Gunther Doehlemann

## Abstract

The maize smut fungi *Ustilago maydis* and *Sporisorium reilianum* are closely related and have similar genomes in terms of size and synteny. While *U. maydis* induces tumors locally at sites of infection, *S. reilianum* systemically colonizes the host and causes symptoms in the inflorescences. To investigate the genetic basis of these differences, an interspecific recombinant hybrid (rUSH) with the mating type system of *S. reilianum* was generated. rUSH exhibited extensive *in-planta* proliferation, showing a *S. reilianum*-like phenotype at all developmental stages except teliospore formation. Transcriptome profiling revealed that expression of pathogenicity-related effector gene orthologs was induced in rUSH, but not in a wild-type hybrid control. Multiple transcriptome comparisons identified 253 differentially expressed one-to-one effector orthologs with distinct regulatory patterns, including cis-, trans-, and rUSH-specific regulation. Functional analysis via CRISPR/Cas9 mutagenesis uncovered three novel virulence factors among the rUSH-specific regulated effectors. Ultimately, rUSH facilitated to identify the transcription factor UmHdp2 as key regulator of *U. maydis-*induced tumorigenesis. Together, these findings highlight the utility of a recombinant, interspecific hybrid in unraveling the molecular mechanisms underlying pathogenic differences in closely related fungal pathogens.

## Introduction

Interspecific hybridization events are a phenomenon that gives rise to the emergence of novel fungal pathogens threatening food sustainability. It enables the exchange of fungal genetic material between species through sexual or parasexual mating (Steensels *et al*., 2021). Fungal hybrids must navigate potential incompatibilities that arise from the evolution of the parental species (Hovhannisyan *et al*., 2020). Before two species can successfully hybridize, they need to overcome pre-mating and post-mating barriers (Steensels *et al*., 2021). The fitness level of new hybrid species is typically influenced by a number of factors, including genomic incompatibilities of separately evolved alleles, competition with parental species, and the characteristics of the ecological environment in which they are situated (Stukenbrock, 2016). Nevertheless, due to pre- and post-mating incompatibilities natural fungal hybrids are rare in nature. However, when such rare hybridization event occurs between closely related species, it can offer valuable insights into evolutionary traits and host specificity.

Smut fungi infect around 1,500 plant species, mainly including Poaceae. Most smuts infect their host systemically without evident disease symptoms in the early infection phase, and replace the inflorescences with teliospores. One example of systemic infection is the head smut causal fungus *Sporisorium reilianum*, which infects maize (*S. reilianum f. sp. zeae*) and sorghum (*S. reilianum f. sp. reilianum*). During infection, *S. reilianum* stays close to the vascular bundles until it reaches the cob primordia (Zuo *et al*., 2021), where it causes the development of multiple female inflorescences and the loss of apical dominance (Ghareeb *et al*., 2015). An exception to the typical smut infection process shows the model organism *Ustilago maydis*, which causes large tumors locally at all aerial infected parts of maize (Brefort *et al*., 2009). The pathogenicity of smut fungi is tightly linked with their sexual cycle, determined by the highly conserved mating type loci. *U. maydis* and *S. reilianum* comprise a tetrapolar mating type system, consisting of two unlinked mating type loci, *a* and *b*. While the *a* locus encodes a pheromone-receptor-system for the recognition of compatible haploid sporidia, the multiallelic *b* locus encodes for the transcription factors (TFs) bEast (bE) and bWest (bW), which act as a heterodimeric key regulator for pathogenicity (Gillissen *et al*., 1992; Schulz, 1990). bE/bW tightly regulates a hierarchical TF network by regulating the master regulator Rbf1 (regulator of b-filament; Heimel *et al*., 2010) and the downstream TFs Biz1 (b-dependent zinc finger protein 1) and Hdp2 (homeodomain transcription factor 2), which are important for the regulation of effector gene expression. The deletion of Biz1 and Hdp2 has been reported to result in a substantial decrease in virulence in *U. maydis* (Flor-Parra *et al*., 2006; Lanver *et al*., 2014), indicating the regulation of the initial wave of effectors that are crucial for the early biotrophic development (Lanver *et al*., 2017). Plant-colonizing microbes secrete a plethora of effector proteins in a spatiotemporal manner to manipulate the host and facilitate host colonization. A genome comparison of *S. reilianum* and *U. maydis* revealed a high degree of synteny, with an overall amino acid sequence identity of all predicted proteins of 74.2% and 62% for secreted proteins, suggesting a more rapid evolution of putative effector genes (Schirawski *et al*., 2010). The genome data of *U. maydis* (Kämper *et al*., 2006) and *S. reilianum* (Schirawski *et al*., 2010) formed the foundation for the analysis of effector orthologs between the two species. Many effector genes are found in gene clusters. Cluster 19A is the largest cluster identified in both species and contains 24 and 29 secreted effector proteins for *U. maydis* and *S. reilianum*, respectively (Brefort *et al*., 2014; Ghareeb *et al*., 2019). 19A deletion mutants of *U. maydis* exhibit an inability to induce the formation of large tumors and failed to develop teliospores (Kämper *et al*., 2006), in line with the absence of teliospore formation in 19A1A2 deletion mutant in *S. reilianum* (Ghareeb *et al*., 2011). A number of effectors have already been characterized in *U. maydis* and *S. reilianum*. These have been found to be either functionally conserved, i.e., as evidenced by the effectors See1 (Redkar *et al*., 2015; Shi *et al*., 2023), Pep1 (Hemetsberger *et al*., 2015), ApB73 (Stirnberg & Djamei, 2016) and Rsp3 (Ma *et al*., 2018) or to have different functions, as exemplified by the effectors Tin2 and Sts2 (Tanaka *et al*., 2019; Zuo *et al*., 2023), respectively. A cross-species transcriptome analysis of *S. reilianum* and *U. maydis* revealed 207 of 336 differentially expressed one-to-one effector orthologs during colonization, suggesting that these genes may contribute to the different disease progressions observed in the two smuts. The findings of this study suggests that the diversification of orthologous effectors in closely related smut fungi can be attributed to two primary factors: transcriptional regulation of effector genes and functional diversification of effector proteins (Zuo *et al*., 2021).

It has been shown previously that the haploid *U. maydis* strain 521 (*Uma1b1*) was able to induce filaments with the haploid strain SRZ2 of *S. reilianum*. However, this induction was observed to occur with a lesser degree of effectiveness in comparison to the compatible *U. maydis* mating partner 518 (*Uma2b2*). This finding suggests that interspecific mating also induces pheromone signaling, indicating the presence of similar molecular networks between the species (Kellner *et al*., 2011). Nevertheless, the complementation of the bE and bW proteins in *U. maydis* with *S. reilianum* resulted in only partial restoration of tumor formation, which indicates a partial functional conservation of the b proteins (Schirawski *et al*., 2005). *S. reilianum* and *U. maydis* infect the same host, comprise similar genomes but no natural hybridization events have been reported so far. However, under laboratory conditions a fungal hybrid of the two smut fungi *Ustilago bromivora* infecting *Brachypodium spp.* and *Ustilago hordei* infecting barley is able to infect *Brachypodium spp.*, while *U. hordei* alone was unable (Bosch *et al*., 2019). Moreover, a fungal hybrid of *S. reilianum* and *U. maydis* was employed to investigate their differences in the disease symptoms. However, after successful formation of a dikaryotic hyphae, a lack of fungal proliferation and a delayed development of disease symptoms was observed for the hybrid compared to the parental species (Storfie & Saville, 2021). Therefore, we hypothesized that the utilization of the mating type system of a single species to generate a recombinant interspecific hybrid of *S. reilianum* and *U. maydis* may enhance the fitness of the hybrid *in planta*, promote increased compatibility of downstream processes, and offer insights into the differential progression of diseases.

In this study, we generated a *U. maydis* strain harboring the *a* and *b* mating loci from *S. reilianum* and developed a recombinant *U. maydis* X *S. reilianum* hybrid (rUSH) as a tool to elucidate differences in the disease development of the two pathogens. rUSH propagated during infection and led to a disease symptom that bear resemblance to *S. reilianum*. Using RNA sequencing, we identified a total of 253 effector orthologs changing their expression patterns in rUSH compared to that between parental species, which led to the identification of novel *U. maydis* and *S. reilianum* virulence factors contributing to the different disease phenotypes. This identified distinct regulation patterns of effector genes: *cis*-, *trans*- and rUSH-specific. Ultimately, overexpressing the TF *Um*Hdp2 was found to enable rUSH to induce maize tumorigenesis which is linked with the upregulation of a set of 41 *U. maydis* effector genes.

## Results

### Generation of the recombinant smut hybrid rUSH

In a two-step sequential transformation, we exchanged the mating type loci *a1* and *b1* in the *U. maydis* haploid wild type strain FB1 against *S. reilianum a1* and *b1*. Using CRISPR-assisted homologous recombination, this resulted in strain FB1_*Sra1b1*. To generate an interspecific hybrid, FB1_*Sra1b1* was used together with the mating compatible partner of *S. reilianum* SRZ2 (*Sra2b2*) and the resulted hybrid was further named rUSH (recombinant *U. maydis* X *S. reilianum* hybrid; Figure 1A). We further exchanged *Sra1* and *Srb1* mating type loci in SRZ1 against *Uma1* and *Umb1* (SRZ1_*Uma1b1*, Figure S1A+B) and used it together with FB2 (rSUH) in infection assays. In contrast to rUSH, rSUH did not reveal a viable hybrid when infected with the haploid *U. maydis* strain FB2. Biomass quantification and WGA-AF488 staining of rSUH-infected maize leaves was performed, which revealed no increase and clump-formation, respectively (Figure S1C+D). This is in line with mating assays by Kellner *et al*. (2011), where 521 (*Uma1b1*)xSRZ2 were able to induce filament formation, while SRZ1x518 (*Uma2b2*) were not. Consequently, rUSH (FB1_*Sra1b1*xSRZ2) was employed for the subsequent analysis of this study.

**Figure 1:**
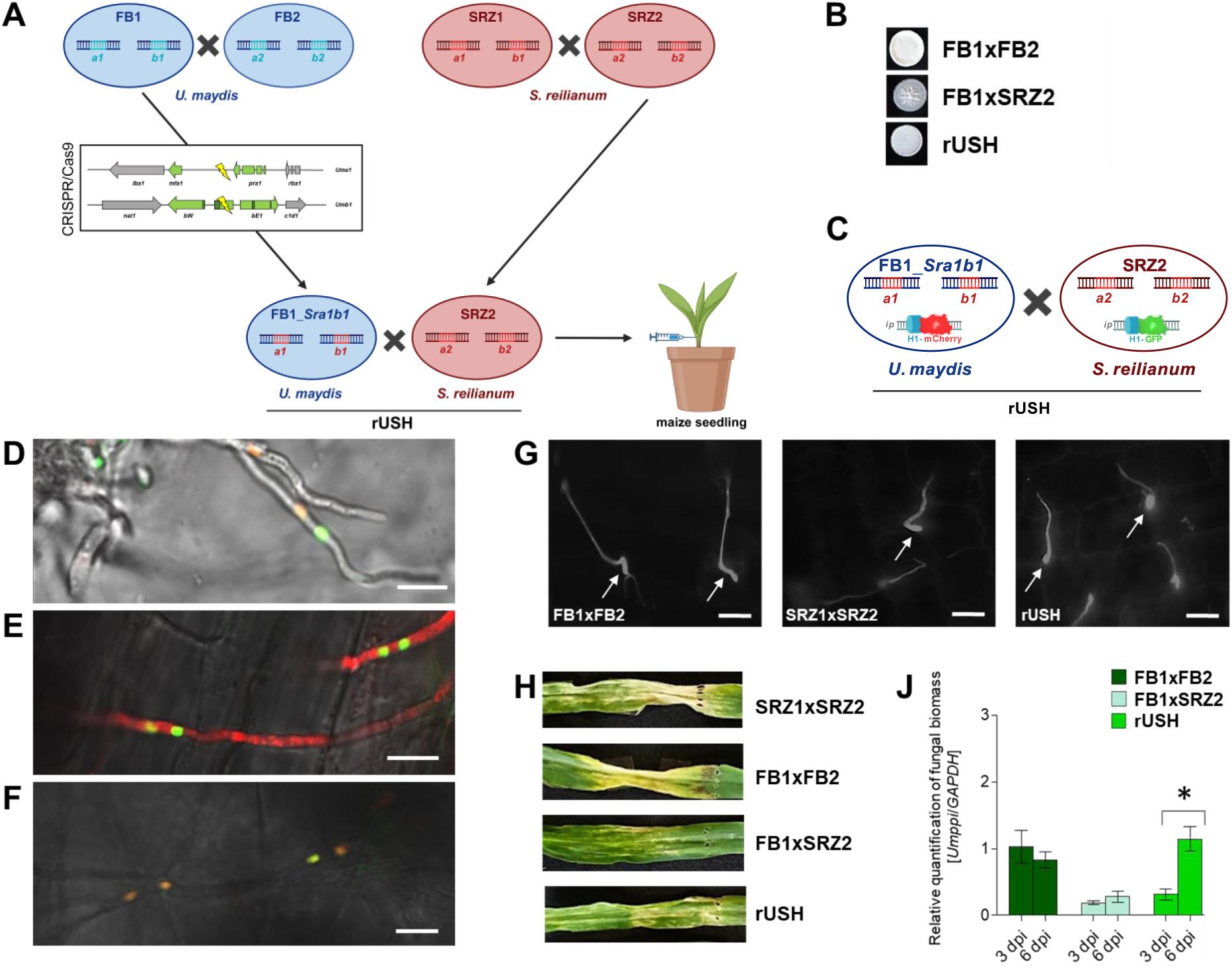
*In planta* proliferation of the recombinant *U. maydis* X *S. reilianum* hybrid (rUSH). **(A)** Generation of rUSH. The mating loci *a1* and *b1* of *U. maydis* FB1 strain were replaced by using CRISPR/Cas9-mediated homologous recombination to generate a mutant containing the *S. reilianum* mating loci *Sra1b1* (FB1_*Sra1b1*). This strain was mixed together with the mating compatible *S. reilianum* strain (SRZ2) to generate rUSH for infection. **(B)** The mix of different mating strains (FB1xFB2, FB1xSRZ2, FB1_*Sra1b1*xSRZ2 (rUSH) on PD-charcoal plate. **(C)** mCherry and GFP-tagged Histon1 were integrated into *ip* locus of FB1_*Sra1b1* and SRZ2, respectively, to visualize the source of the nuclei. **(D)** Confocal microscopy of FB1_*Sra1b1*_H1_mCherryxSRZ2_H1_GFP on activated charcoal. **(E)** Confocal microscopy of 2 dpi maize leaf infected with FB1_*Sra1b1*_mCherry(cytosolic)xSRZ2_H1_GFP. **(F)** Confocal microscopy of 2 dpi maize leaf infected with FB1_*Sra1b1*_H1_mCherryxSRZ2_H1_GFP. **(G)** Calcofluor white staining of maize leaves (20-22 hpi) shows the formation of the appressoria-like structures (indicated in white arrows). **(H)** Phenotype (6 dpi) of SRZ1xSRZ2-, FB1xFB2-, and rUSH-infected maize leaves. **(J)** Quantification of fungal biomass at 3 dpi and 6 dpi. Housekeeping genes *ZmGAPDH* and *Umppi* were used for quantification. Significance was calculated by Students t-test: *, p<0.05. Scale bar (D-G): 10 µM.

### rUSH proliferates *in planta* and reveals a *S. reilianum*-like phenotype

To investigate the viability of rUSH, we tested its ability to form filaments. Therefore, liquid culture of rUSH was dropped on PDA plates supplemented with activated charcoal, which revealed a typical fuzzy filament (Day & Anagnostakis, 1971; Figure 1B). To confirm the formation of a dikaryotic filament, an mCherry- and GFP-tagged Histon1 (H1) were integrated into the *ip* locus of FB1_*Sra1b1* and SRZ2 (Figure 1C), respectively. These strains resulted in rUSH with each one red- and one green-fluorescent nucleus, deriving from *U. maydis* and *S. reilianum*, respectively. Confocal microscopy of the filament from PDA+charcoal (Figure 1F) and *in-planta* 2 days post infection (dpi) verified the presence of one of each mCherry- and GFP- labeled nucleolus in one filament cell (Figure 1D). As a control, an FB1_*Sra1b1* strain with a cytosolic (pro*^actin^*) mcherry was used together with SRZ2 (Figure 1E). Following the life cycle of *U. maydis* further, we investigated the ability of rUSH to form penetration hyphae. Calcofluor white staining of 22 hpi (hours post infection) infected maize seedlings resulted in an observation of appressoria-like structures for *U. maydis*, *S. reilianum*, and rUSH (Figure 1G). In summary, rUSH was found to be capable to form dikaryotic filaments and appressoria-like structures.

Next, the infection phenotypes of rUSH were assessed in comparison to *U. maydis* and *S. reilianum*. Maize seedlings (cultivar: Golden Bantam) infected with rUSH, *U. maydis* and *S. reilianum* were assessed at 6 dpi (Figure 1H). Seedling infections were monitored at 3 weeks post infection (wpi) and 7 wpi maize plants (cultivar: Gaspe Flint), respectively (Figure 2). Interestingly, rUSH caused a *S. reilianum*-like phenotype with the induction of chlorotic and necrotic spots at 6 dpi and at 7 wpi, a few infected plants show leafy tassels and ears (Figure 2). In general, the rUSH phenotypes were similar but milder compared to *S. reilianum* in all infection experiments performed in this study. We measured the relative fungal biomass of maize leaves infected with FB1xFB2, FB1xSRZ2 and rUSH at 3 dpi and 6 dpi. rUSH revealed an increase in biomass between 3 dpi and 6 dpi, in contrast to the control FB1xSRZ2 (Figure 1J). To gain further insight into the underlying gene expression patterns, we conducted an RNA-seq experiment at 3 dpi. In the wild type FB1xSRZ2 hybrid, expression of known crucial effector genes including *cmu1* (Djamei *et al*., 2011), *pit2* (Misas Villamil *et al*., 2019; Mueller *et al*., 2013), *stp2*, *stp3*, *stp5* and *stp6* (Ludwig *et al*., 2021) was not detected (Figure S3B), which most likely explains the inability to further proliferate in the host tissue. In contrast, we observed significant induction of 119 *U. maydis* and 294 *S. reilianum* effector one-to-one orthologs (Zuo *et al*., 2021) in rUSH (Figure S4). This effector gene induction in rUSH can be expected to result in an efficient suppression of plant defense responses. Consequently, the recombinant hybrid was able to maintain the biotrophic interaction, which is reflected by the increasing fungal biomass from 3 dpi to 6 dpi (Figure 1J). Together, this suggests that the reconstruction of a single species’ mating type system leads to a stable and compatible downstream signaling in the interspecies hybrid, and consequently to a successful host infection.

**Figure 2:**
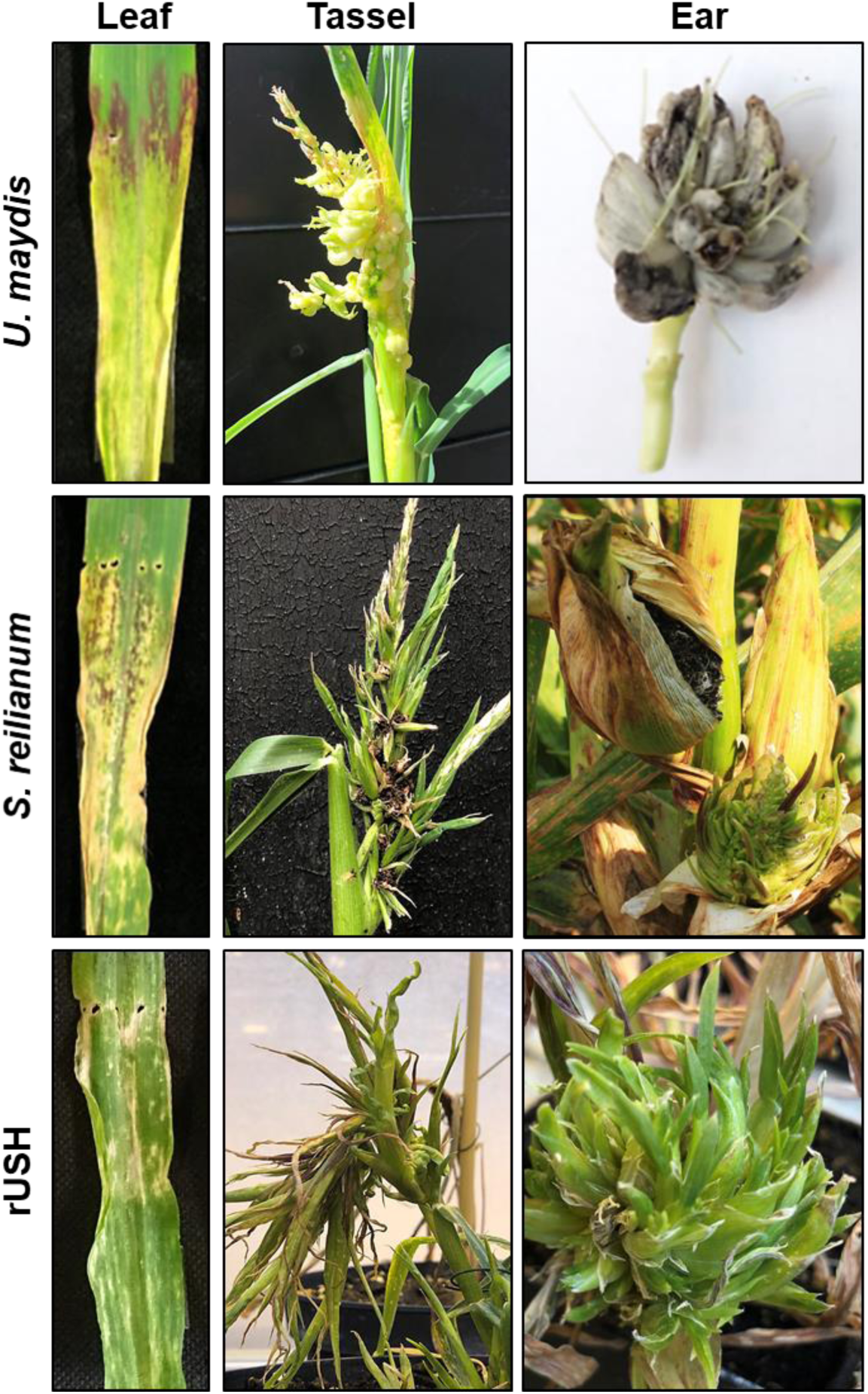
rUSH shows a *S. reilianum*-like phenotype on different maize organs. Phenotype of *U. maydis* wild type FB1xFB2, *S. reilianum* wild type SRZ1xSRZ2 and rUSH at 6 dpi (Leaf), 3 wpi *U. maydis* (Tassel) or 7 wpi for *S. reilianum* (Tassel, ear) and *U. maydis* (Ear).

### *S. reilianum* effector gene orthologs are dominantly expressed in rUSH

To understand how effector ortholog expression contributes to the observed *S. reilianum*-like phenotype, qRT-PCR of two effector genes *pit2* and *see1* (Mueller *et al*., 2013; Redkar, *et al*., 2015) was performed for *S. reilianum*-, *U. maydis*-, and rUSH-infected maize seedlings at 20-22 hpi, 3 dpi and 6 dpi (Figure S2A). For the *U. maydis* orthologs *Umpit2* and *Umsee1*, the expression levels were significantly reduced in rUSH at 6 dpi compared to FB1xFB2. In contrast, the *S. reilianum* orthologs *Srpit2* and *Srsee1* were significantly higher expressed in rUSH at 6 dpi compared to *S. reilianum*, indicating an opposite expression pattern of these orthologs in rUSH (Figure S2A). qRT-PCR results suggested an in general lower expression of *U. maydis* orthologs compared to the *S. reilianum* genes. To test whether the RNAi machinery in *S. reilianum*, which is absent in *U. maydis* (Schirawski *et al*., 2010), is associated with this downregulation of the *U. maydis* orthologs in rUSH, a dicer (*sr16838*) knock-out was generated in the SRZ2 background (Figure S2B). However, when the gene expression levels of *Umsee1, Umtin2* and *Umnlt1* were measured in rUSH and two independent rUSH dicer deletion strains, no change in expression levels was observed, suggesting that the observed downregulation effect of *U. maydis* orthologs is not caused by RNAi (Figure S2D).

Next, we tested whether a nucleus-specific suppression of the *U. maydis* gene expression takes place. To this, we used the effector gene *Umtin2*, which had been shown to induce maize anthocyanin formation (Tanaka *et al*., 2014). Moreover, it’s ortholog SrTin2 can neither complement the virulence, nor the anthocyanin formation of the mutant Δ*Umtin2* (Tanaka et al., 2019). Thus, the observation that rUSH infection did not induce anthocyanin formation, is in accordance with the ortholog-specific gene expressions of *Srtin2* in rUSH (Figure 3A+B). To investigate whether a locus-specific inhibition of *Umtin2* expression takes place in *U. maydis*, we used the promoter of *Srtin2*, pro*^Srtin2^* to control the expression of *Umtin2* in the native *Umtin2* locus of FB1_*Sra1b1* (Figure 3C). As a control, we replaced the coding sequence of *Srtin2* against *Umtin2* in its native locus in SRZ2 (Figure 3D) by using CRISPR-mediated knock-in. In an infection assay, FB1_*Sra1b1*_pro*^Srtin2^*_*Umtin2*xSRZ2 and FB1_*Sra1b1*xSRZ2_pro*^Srtin2^*_*Umtin2* induced anthocyanin formation, which is absent in rUSH (Figure 3D). This demonstrates that the induction of *Umtin2* in rUSH can be restored by maintaining the *cis*-regulatory element of the promoter of *Srtin2*, which also excludes a locus-specific suppression of the *U. maydis* ortholog. However, at present regulatory differences between *Umtin2* and *Srtin2* are far away from being fully understood on a genome-wide scale, and this might either involve variations in the *cis*-regulatory elements, or different TFs between the species.

**Figure 3:**
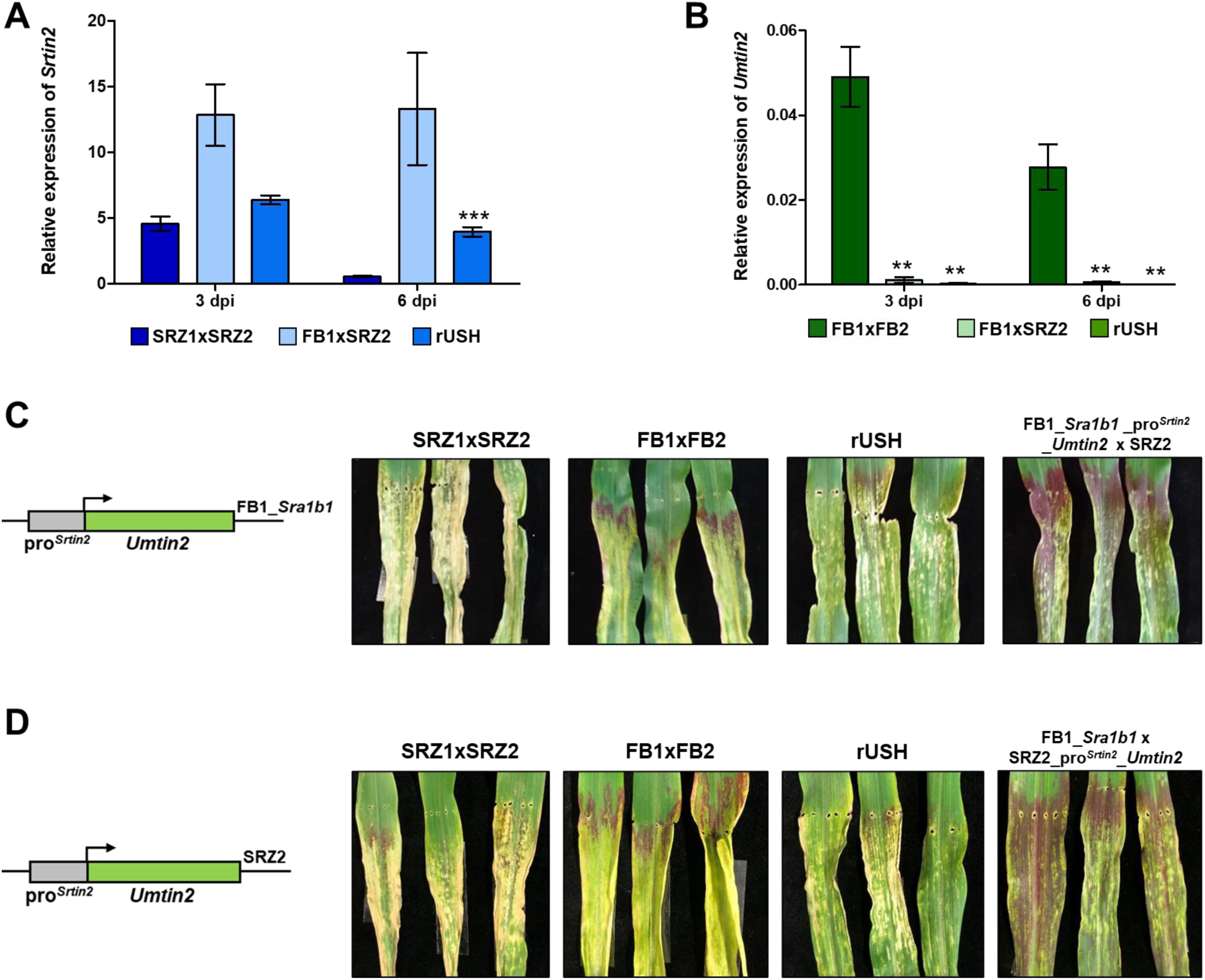
The differential regulation of *Umtin2* ortholog in rUSH is caused by the *cis*-regulation in the promoter. **(A)** qRT-PCR detection of *Srtin2* expression in SRZ1xSRZ2, FB1xSRZ2, and rUSH at 3 and 6 dpi. *Srppi* is used for normalization. **(B)** Relative expression of Umtin2 in FB1xFB2, FB1xSRZ2 and rUSH. *Umppi* is used for normalization. **(C,D)** Phenotype assessment of rUSH: pro*^Srtin2^* controlling *Umtin2* in FB1_*Sra1b1* and **(C)** pro*^Srtin2^* controlling *Umtin2* in *S. reilianum* SRZ2. **(D)** Photos of typical phenotypes of different infections at 6 dpi. Error bars (standard deviation) were calculated from three biological replicates. Significant differences were calculated based on Students’ t-test: **, p<0.01; ***, p<0.001).

### Transcriptome analysis reveals distinct expression patterns of one-to-one effector genes in rUSH

For a genome-wide profiling of effector ortholog expression in rUSH, RNA-sequencing of *U. maydis* (FB1xFB2), *S. reilianum* (SRZ1xSRZ2) and rUSH was conducted at 20-22 hpi (penetration stage), 3 dpi (early biotrophic development) and 6 dpi (*U. maydis* tumorigenesis) (Figure 4A). In line with previous findings, where relative reads were compared between species (Zuo *et al*., 2021), we obtained significantly more reads for *S. reilianum* at 3 and 6 dpi as compared to *U. maydis* (Figure 4B). In rUSH, the read contribution from each fungal species was comparable at 20 hpi and 3 dpi, and slightly more reads were obtained from *U. maydis* at 6 dpi (Figure 4C). Epidermal penetration and initiation of the biotrophic development (20-22 hpi) is similar in both pathogens (Zuo *et al*., 2021). During leaf colonization the development of the two pathogens gets increasingly distinct, particular at 6 dpi, when only *U. maydis* but not *S. reilianum* is inducing tumor formation. This corresponds with an increasing differential expression of effector orthologs over time: we identified 75 differentially expressed one-to-one effector genes (log_2_FC>1 or <-1; p<0.05) at 20-22 hpi, 114 at 3 dpi, and 152 at 6 dpi (Figure 4D+E). Based on the expression profiles of the one-to-one orthologs between WT and rUSH, ortholog clustering was performed, which revealed 8 clusters with distinct expression patterns (Figure 4F). After examining the expression levels of one-to-one orthologs in *U. maydis*, *S. reilianum*, and rUSH, we have identified three distinct expression patterns: *cis* (84), *trans* (61), and rUSH-specific (108) and these patterns can be associated with the different clusters observed. *Cis*-regulated expression reflects the same trend of differential expression between *U. maydis* and *S. reilianum* and within rUSH, indicating *cis*-regulatory elements in the promoter region contributed to the differential expression. *Trans*-regulated expression indicates the differential expression between *U. maydis* and *S. reilianum* which is gone in rUSH, influenced by the presence/absence of a TF or the different threshold of TF expression. Lastly, the rUSH-specific expression showed two different observations: reverse expression (58), in which one ortholog shows a dominant expression in either *U. maydis* or *S. reilianum*, while in rUSH the opposite ortholog has higher expression, and ortholog-specific expression (50), where only one ortholog changes the expression in rUSH compared to *U. maydis* and *S. reilianum* (Figure 5A). Within the rUSH-specific reverse expression pattern, previously characterized effector genes are included, i.e. *pit2*, *tip1*, *tip2*, *tip4-6* (Bindics *et al*., 2022; Huang *et al*., 2024, see Table S1), which are known to contribute to *U. maydis* virulence in maize leaves.

**Figure 4:**
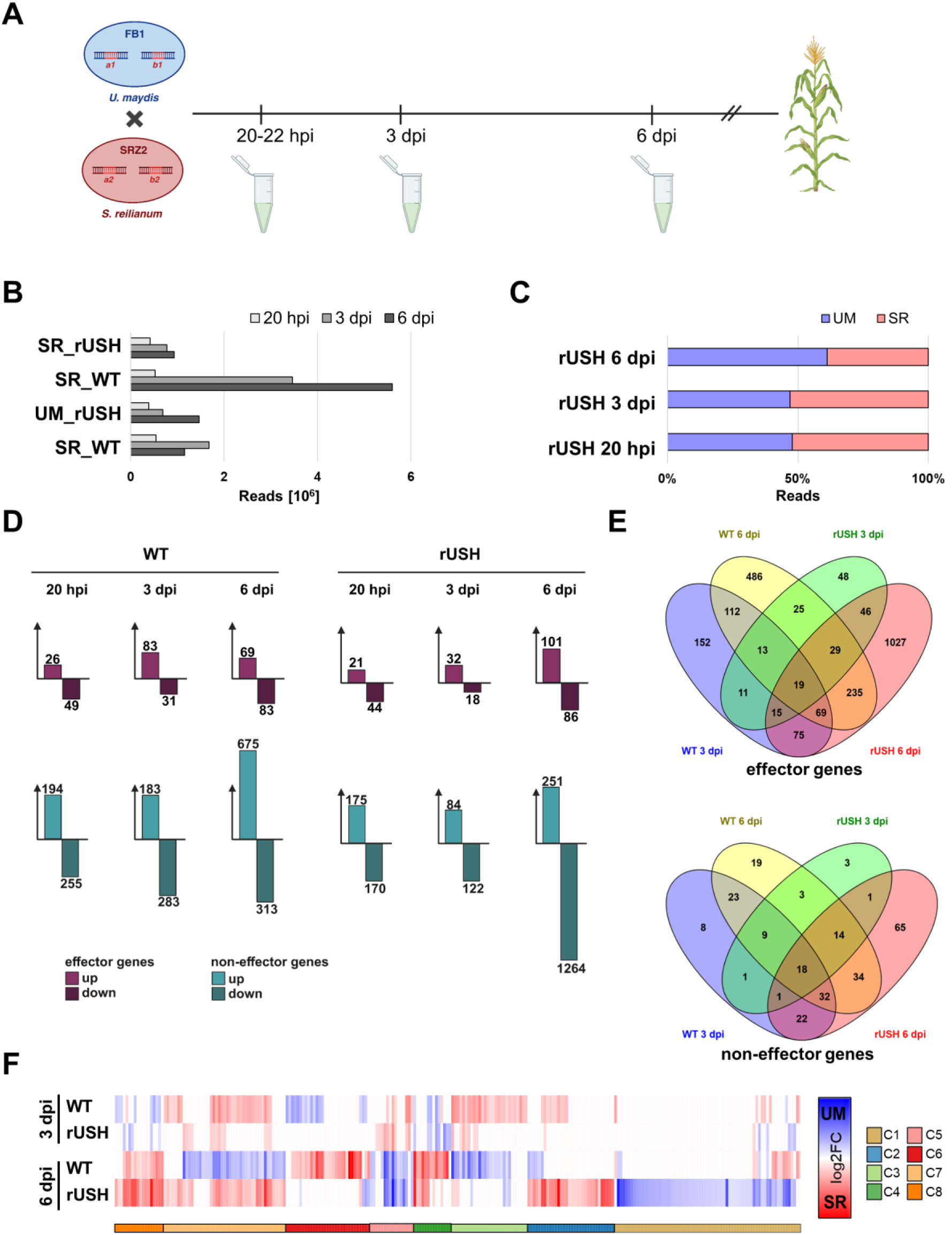
RNA-seq of rUSH compared to FB1xFB2 and SRZ1xSRZ2 at 20-22 hpi, 3 dpi, and 6 dpi. **(A)** Time points of RNA-seq sampling. At 20-22 hpi, fungal material on the plant surface was enriched using liquid latex. At 3 dpi and 6 dpi, 4 cm of plant material was used for the extraction of total RNA. **(B)** Total reads were obtained from each species at 20 hpi, 3 dpi, and 6 dpi. **(C)** Relative reads from *U. maydis* and *S. reilianum* in rUSH at 20 hpi, 3 dpi and 6 dpi. **(D)** Bar plots show the number of differentially expressed one-to-one effector orthologs and non-effector orthologs (Zuo *et al*., 2021) between *S. reilianum* and *U. maydis* (WT) and between *S. reilianum* and *U. maydis* within rUSH. **(E)** Venn diagrams show the overlap of effector orthologs and non-effector orthologs at 3 dpi and 6 dpi in FB1xFB2 (WT) and rUSH. **(F)** Heatmap shows the log_2_ fold change (log_2_FC) between *U. maydis* and *S. reilianum* orthologs between wildtype or within rUSH. Ortholog clustering was performed based on the expression profiles between WT and rUSH using one minus pearson correlation and KMeans clustering.

**Figure 5:**
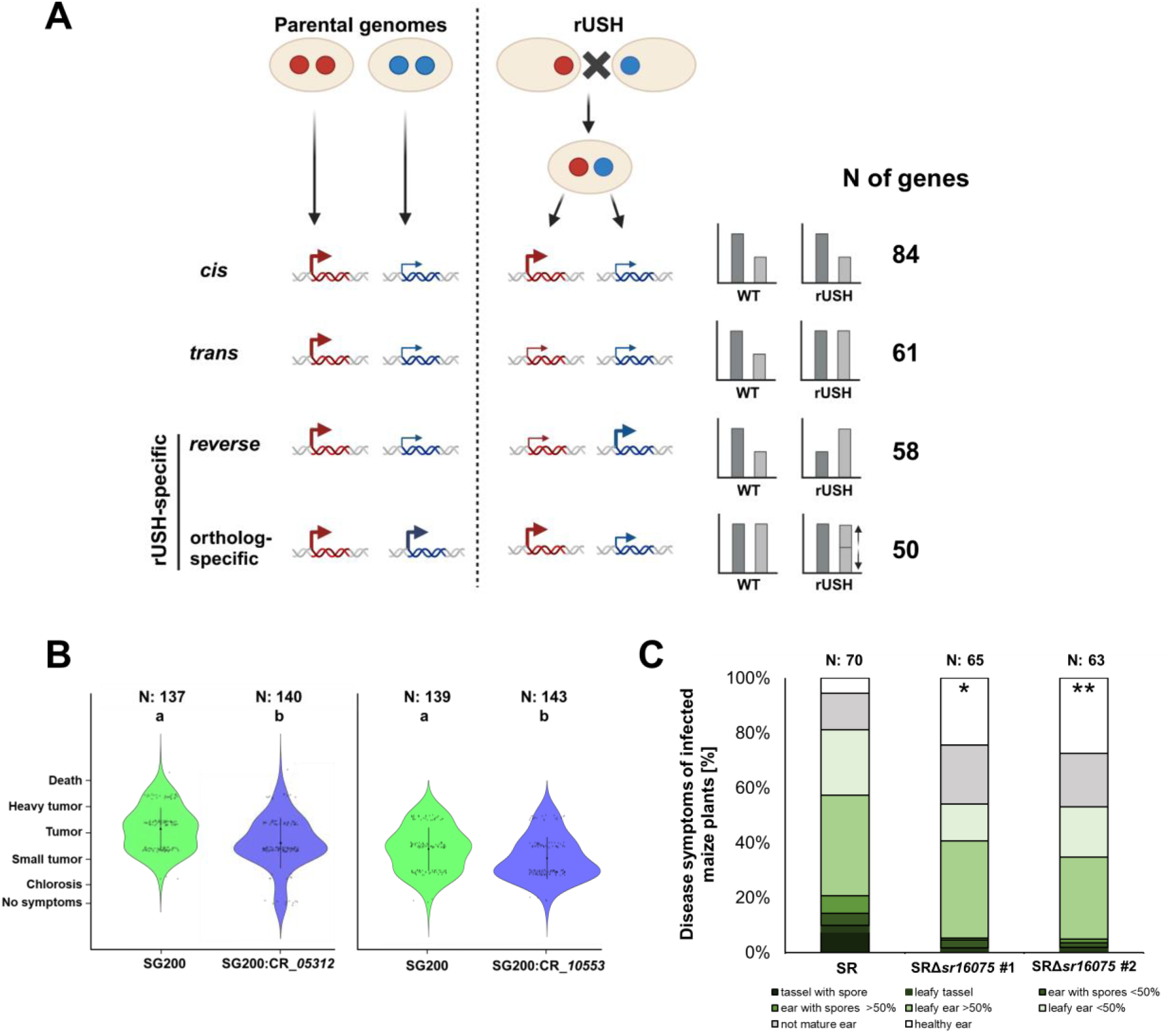
Effector orthologs in rUSH reveal novel expression patterns compared to wild type. **(A)** Expression patterns of differentially expressed one-to-one effector orthologs in rUSH. Effector orthologs between the wild type *U. maydis* and *S. reilianum* are differentially expressed: i) *Cis*: remains the same expression in rUSH, ii) *Trans*: exhibits an equal expression between the two orthologs within rUSH. iii) Reverse expression: shows the opposite expression in rUSH, iv) Ortholog-specific expression: only one ortholog changes the expression within rUSH. The effector gene expression was calculated by dividing the *S. reilianum* transcripts per million (TPM) by *U. maydis* TPM of the wild types FB1xFB2 and SRZ1xSRZ2 (SR_WT/UM_WT) as well as within rUSH (SR_rUSH/UM_rUSH). **(B)** Infection assay of SG200, SG200:CR_UMAG_*05312* and SG200:CR_*UMAG_10553* at 12 dpi. 7-days old maize seedlings were infected with SG200 and mutant strains. Significant differences were calculated based on Tukey’s two-way ANOVA. **(C)** Infection assay of early flowering maize cultivar Gaspe Flint with *S. reilianum* wild type SR (SRZ1xSRZ2) and two independent mutants of *sr16075* (SRZ1Δ*sr16075*xSRZ2Δ*sr16075*: SRΔ*sr16075* #1 and #2). 7-days old maize seedlings were infected with a final OD of 1 of the mating type combinations. Disease symptoms were scored 8 wpi as described previously (Ghareeb *et al*., 2019).

### Effector genes with rUSH-specific expression encode virulence factors

Effector genes that are typically highly induced during *U. maydis* infection were significantly downregulated in rUSH. This finding is of particular interest with regard to the absence of tumor induction by rUSH. In line with this *S. reilianum*-like phenotype, we hypothesized that those *U. maydis* effector genes that are downregulated in rUSH are likely linked to tumorigenesis. To follow this assumption and to identify effector genes that contribute to *U. maydis*-induced tumorigenesis, we set a stringent cut-off of an at least 5-fold higher expression in the *U. maydis* wild type compared to rUSH. The resulting candidate list of 14 tumorigenic effector candidate genes (Table S1) includes effector genes that have already been functionally characterized, including the *U. maydis* virulence factors Pep1, Sts2, Tip6, and Tip7 (Doehlemann *et al*., 2009; Huang *et al*., 2024; Khan *et al*., 2024; Zuo *et al*., 2023). To identify novel virulence factors, six candidate genes that had not been investigated in previous studies were selected to further functional characterization (Table S1). CRISPR/Cas9 (CR) frameshift mutants were generated in *U. maydis* and the resulting mutant strains were tested in maize infection assays. Strikingly, two of the selected candidate genes (*UMAG_05312* and *UMAG_10553*) were identified as virulence factors, i.e. the respective mutant strains showed significantly reduced tumorigenesis compared to the progenitor strain SG200 (Figure 5B). This result confirms the hypothesis that reverse-expressed effectors with downregulation in rUSH are involved in tumor formation.

In a complementary approach, we also checked the expression levels of *S. reilianum* effector genes in rUSH compared to *S. reilianum*. This revealed 108 *S. reilianum* effector genes in rUSH with at least 2-fold higher expression levels compared to *S. reilianum* (Table S1). Using CRISPR/Cas9 mutagenesis, we generated compatible *S. reilianum* knock-out strains of the most highly expressed effector candidate *sr16075* in our list. Maize infection assays revealed a reduction of virulence with a significant increase in healthy ears compared to *S. reilianum* wild type at 7 wpi (Figure 5C). Thus, mutation of effector genes with rUSH-specific expression pattern in rUSH identified novel virulence factors in both *U. maydis* and *S. reilianum*.

### The transcription factor Hdp2 regulates tumorigenic effector genes

The differential expression patterns of *U. maydis* / *S. reilianum* wild type strains and rUSH might be caused either by different *cis*-regulatory elements in the promoter region of the effector genes, or by the presence/absence or a certain threshold of a TF. To identify TFs implicated in the transcriptional regulation of differential gene expression and, consequently, the different disease progression of *U. maydis* and *S. reilianum*, we sought out one-to-one non-effector genes with a predicted DNA-binding function. This led to the identification of 78 candidate TFs (Table S1). We evaluated the candidates with a focus on TFs with lower *U. maydis* ortholog expression in rUSH compared to the wild type. Furthermore, we selected reverse-expressed TF candidates with a high expression of the *S. reilianum* one-to-one ortholog in rUSH, while in the *U. maydis*-*S. reilianum* comparison the *U. maydis* ortholog is higher expressed. Based on these parameters, five candidates were selected to generate CR frameshift mutants in the *U. maydis* background for subsequent pathogenicity assays (Figure S4F). However, none of the five candidate genes exhibited a role in *U. maydis* virulence (Figure S4A-E). We therefore compared the expression patterns of known, pathogenicity-related *U. maydis* TFs in rUSH with their expression in *U. maydis.* Here we observed an overall higher expression of their *S. reilianum* one-to-one orthologs in rUSH (Figure S4G). Based on this observation we hypothesized that this low expression of *U. maydis* TF-orthologs might be functionally linked with the *S. reilianum*-like phenotype of rUSH. If true, one would therefore expect that an overexpression of these TFs in rUSH might shift the pathogenic behavior of the hybrid towards *U. maydis*, i.e. the formation of leaf tumors. Support of this hypothesis is also given by a previous study, where overexpression of *Umrbf1* and *Umhdp2* in 521xSRZ2 revealed the induction of small tumors in a rare event (Storfie & Saville, 2021). We therefore overexpressed the *U. maydis* TFs *Umrbf1*, *Umnlt1*, *Umfox1*, *Umros1* and *Umhdp2* (Heimel *et al*., 2010; Lanver *et al*., 2014; Lin *et al*., 2021; Tollot *et al*., 2016; Zahiri *et al*., 2010) in rUSH, using the promoter of *U. maydis* effector gene *cmu1* (*pro^Umcmu1^*; Djamei *et al*., 2011) that confers high expression in *U. maydis* throughout the biotrophic development (Djamei *et al*., 2011; Lanver *et al*., 2018). The overexpression of *Umrbf1*, *Umnlt1*, *Umfox1*, *Umros1* in rUSH did not cause a significant change in rUSH infection (Figure S4H). However, when rUSH overexpressing *Umhdp2* was infected to maize leaves, tumor formation locally at sites of infection was observed (Figure 6A) in all tested plants, which strikingly resembled the typical disease phenotype caused by *U. maydis*.

**Figure 6:**
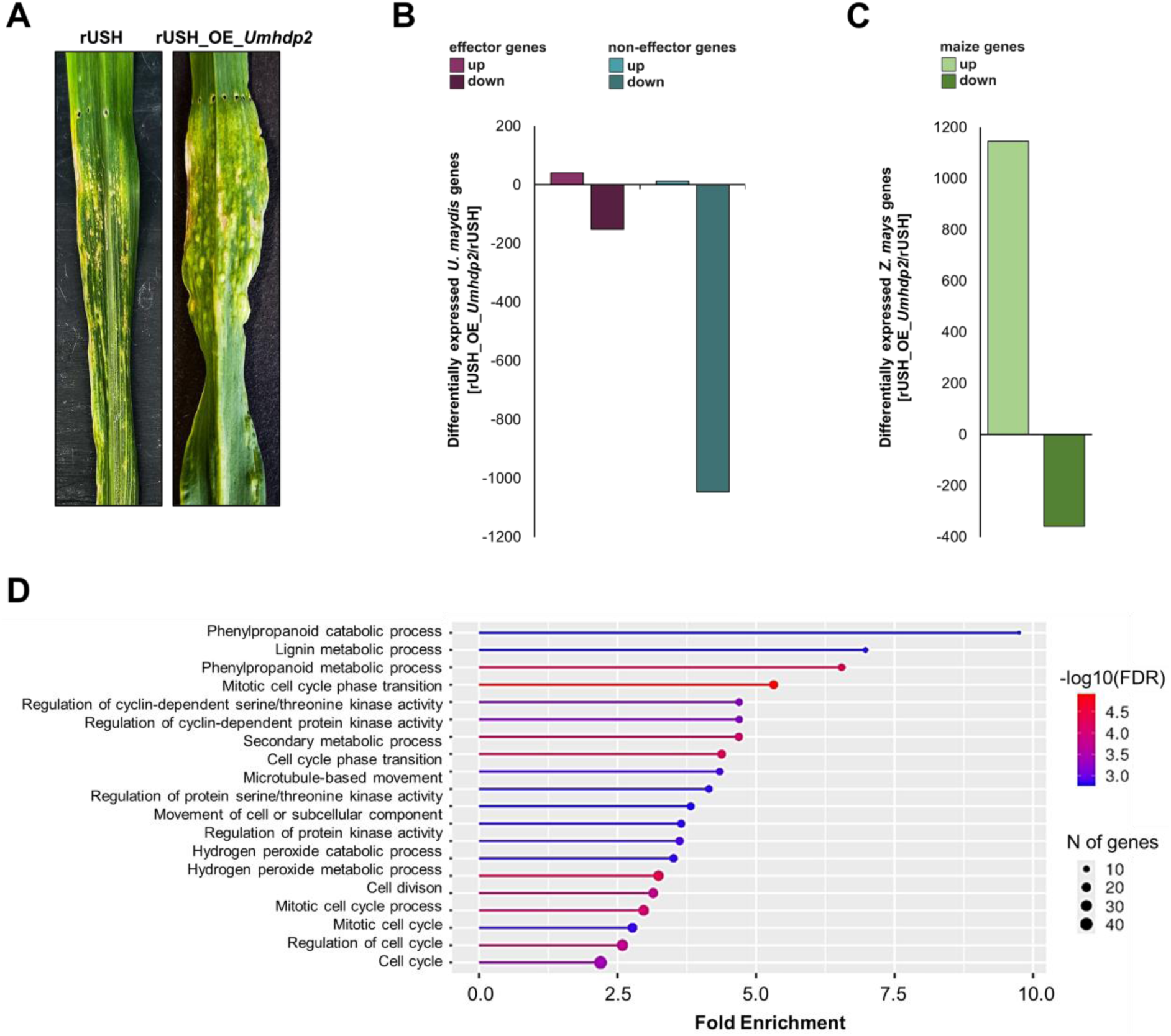
Overexpression of the transcription factor *Umhdp2* leads to tumor formation on rUSH-infected maize leaves. **(A)** Phenotype of rUSH and rUSH_OE_*Umhdp2* at 6 dpi. **(B)** Differentially expressed *U. maydis* effector genes and non-effector genes. **(C)** Differentially expressed maize genes in rUSH_OE_*Umhdp2* in comparison to rUSH (rUSH_OE_*Umhdp2*/rUSH). **(D)** GO Enrichment analysis of 1,145 upregulated maize genes in rUSH_OE_*Umhdp2* compared to rUSH (shinyGo 0.66).

Thus, genes being induced by UmHdp2 in rUSH are likely to be functionally linked with tumorigenesis. To identify the genes, an RNA-sequencing experiment was conducted, comparing rUSH_OE_*Umhdp2* and rUSH. This identified 41 *U. maydis* effector genes being upregulated by *Um*Hdp2, including 13 and 5 genes which reside in the effector clusters 19A and 6A (Kämper *et al*., 2006), respectively. Additionally, 12 non-effector genes were upregulated in rUSH_OE_*Umhdp2* (Figure 6B). In line with the observed tumor formation, we identified 1.145 upregulated maize genes (Figure 6C) in rUSH_OE_*Umhdp2* vs. rUSH. Here, genes associated with metabolic/catabolic processes and cell cycle control were enriched. These processes have previously been found to be associated with *U. maydis*-induced tumor induction (Doehlemann *et al*., 2008; Redkar *et al*., 2015). Within the genes enriched in cell cycle processes, 14 genes were associated to the regulation of mitotic nuclear division, 18 in the regulation of mitotic cycle and 16 in cell cycle transition (Figure 6D). Strikingly, this also included the leaf developmental markers *ZmGIF1* and *ZmSHR1*, which we recently found to be induced by the tumor-inducing effector Sts2 (Zuo *et al*., 2023). The identification of *ZmGIF1* and *ZmSHR1* in two independent RNA-seq analyses as potential targets of *U. maydis* in maize, involved in tumorigenesis (this study; Zuo *et al*., 2023), renders them promising candidates for investigating the underlying mechanisms of *U. maydis*-induced leaf tumor development in the future.

## Discussion

*U. maydis* and *S. reilianum* are closely related pathogen species that infect the same host plant, but they differ fundamentally in their infection styles. Apart from two characterized tumorigenic effectors See1 and Sts2 (Redkar, Hoser, *et al*., 2015; Zuo *et al*., 2023), the genetic basis of *U. maydis* tumor formation remains largely unknown. In this study, we generated a recombinant *U. maydis* X *S. reilianum* hybrid (rUSH) as a tool to investigate the different disease development of the two pathogens, in particular with regard to tumorigenesis. rUSH successfully colonized maize leaves and inflorescences and displayed a *S. reilianum*-like phenotype. Transcriptome analysis of *U. maydis*, *S. reilianum* and rUSH identified rUSH-specific expression patterns of effector orthologs, which revealed novel virulence factors of both pathogens and identified *Um*Hdp2 as key transcriptional regulator for the induction of *U. maydis*’ tumorigenesis.

Previous approaches to generate smut hybrids between *U. maydis* and *S. reilianum* without altering the mating type loci, such as protoplast fusion (Heinze, 2009)or the laboratory hybrid 521xSRZ2 (Storfie & Saville, 2021) resulted in valuable insights but were limited in their pathogenic development. In the Ustilaginaceae, the sexual life cycle is intricately connected to pathogenicity, which is influenced by the compatibility of their mating type systems. The high conservation of the mating type systems between *U. maydis* and *S. reilianum* eliminated pre-mating barriers, facilitating successful mating and the formation of dikaryotic hyphae. Comparative genetics along with sexual compatibility tests conducted by Kellner *et al*. (2011) showed the compatibility between haploid strains of *U. maydis* and *S. reilianum*. In accordance with our observations on rUSH, only *U. maydis* 521 (*Uma1b1*) was observed to induce filaments with the second mating partner of *S. reilianum* (SRZ2) and not *vice versa* (Kellner et al., 2011).

Our transcriptome analysis revealed a lack of expression of key effector genes in wild type FB1xSRZ2 hybrids, which is in line with the attenuated host colonization of 521xSRZ2 (Storfie & Saville, 2021) and FB1xSRZ2 (this study). This reflects the general fitness deficits in the natural combination of interspecies mating types, which is overcome by the recombinant mating type loci in rUSH. Notably, leafy ears, a typical symptom of *S. reilianum*, were observed in rUSH infection at 7 wpi, indicating systemic growth of rUSH up to 7 wpi. However, in our hands rUSH was unable to switch to produce teliospores, which are necessary for further sexual reproduction (Samarasinghe *et al*., 2020). We assume that rUSH is incapable of completing its life cycle due to post-mating incompatibilities, which are associated with signaling processes involved in sexual reproduction. This is also supported by our unsuccessful attempts to restore teliospore formation by overexpressing the *U. maydis* TF *ros1* in rUSH, which was shown to be crucial for karyogamy and matrix formation (Tollot et al., 2016 and data not shown). To overcome post-mating barriers and obtain teliospores in rUSH, future approaches might include overexpression of TFs which are highly induced during the reproductive phase and might be involved in the regulation of developmental genes essential for sporogenesis.

In hybrids, gene expression is significantly influenced by a complex interplay of *cis*-regulatory elements and *trans*-regulatory factors, which are affected by the divergence of the parental species, alterations in chromatin structure and modifications, as well as RNAi (Combes et al., 2015). *U. maydis* lacks an RNA interference (RNAi) machinery, which is a distinctive feature among the smuts (Schirawski *et al*., 2010). The substantial role of the RNAi machinery in the observed downregulation of *U. maydis* effector genes in rUSH was refuted, as dicer gene deletion in *S. reilianum* did not alter expression levels. After the so-called “transcriptomic shock”, which frequently occurs following the merging of different genomes (Steensels *et al*., 2021), RNA-seq of rUSH-infected maize leaves revealed 253 differentially expressed one-to-one effector orthologs, including rUSH-specific expression patterns. This rUSH-specific expression is likely the result of an interaction between *cis*- and *trans*-regulation, which differs from earlier studies on interspecific hybrids where gene expression was either equally or more strongly affected by *cis*-regulation compared to *trans*-regulation (Hill et al., 2021; Runemark et al., 2024). Consistent with our results, hybrid-specific expression has also been observed in yeast hybrids, although to a lesser degree than what was noted in rUSH (Tirosh *et al*., 2009). Furthermore, it has been demonstrated for rUSH that the effector gene *Umtin2* is differentially expressed between the species and within rUSH, which is in line with a previous study (Zuo *et al*., 2021). The variation in *tin2* expression is likely attributable to *cis*-regulatory elements within the promoter region, which play a crucial role in determining the different expression levels and could be different among different species.

Effector gene expression is tightly regulated in a spatiotemporal manner by a hierarchical network of TFs (Feldbrügge et al., 2004; Heimel et al., 2010; Lanver et al., 2018; Skibbe et al., 2010). When *Umhdp2* was overexpressed in rUSH, the formation of leaf tumors was observed in all infected maize seedlings, demonstrating that Hdp2 holds a key role as a regulator of tumorigenesis-associated effector genes. The transcriptome analysis of rUSH_OE_*Umhdp2* vs. rUSH identified a set of 41 tumor-inducing effector genes in *U. maydis*. Using rUSH as a platform, future work, involving consecutive knock-out mutants and in parallel overexpression of tumor-inducing effector genes, will help to unravel a minimal key set of effectors being required for the induction of plant tumors. Taken together, the generation of a recombinant, interspecific hybrid disclosed the regulatory patterns of effector orthologs in *U. maydis* and *S. reilianum*. The characterization of rUSH facilitated the identification of new virulence factors in both *U. maydis* and *S. reilianum* and highlighted maize genes that may play a role in the tumor pathways initiated by *U. maydis* infections. Above all, however, the establishment of rUSH is a decisive step towards the mechanistic elucidation of the genetic basis of fungal tumor formation in plants.

## Material and Methods

### Strains, growth conditions, and plant infections

For the RNA-seq analysis and infection studies, the mating compatible strains used, including *U. maydis* (FB1 and FB2), *S. reilianum* (SRZ1xSRZ2), and the recombinant hybrid (rUSH, FB1_*Sra1b1*xSRZ2). Infection assays for effector candidate gene knock-outs (KOs) were carried out using SRZ1 and SRZ2. CRISPR/Cas9 frameshift mutants and knock-in mutants were generated for *U. maydis* (Schuster *et al*., 2016; Zuo *et al*., 2020) and *S. reilianum* (Werner *et al*., 2024)according to previously established methods. In cases where no donor template was available, CRISPR/Cas9 mutagenesis was implemented as outlined by (Zuo et al., 2021). The DNA sequence of the genomic region of interest was sent for sequencing to verify the presence of a premature stop codon. For strains generated through CRISPR/Cas9-assisted homologous recombination (HR), Southern blot analysis was conducted. All strains were cultivated in YEPS_light_ liquid medium at 28°C with shaking at 200 rpm, or on potato dextrose agar plates (PD). For plasmid cloning the *Escherichia coli* strain Top10 was used, grown in dYT liquid medium or on YT plates, both supplemented with the appropriate antibiotics. For the infection assays, maize plants were grown under regulated conditions: 16 h of light at 28°C and 8 h of darkness at 22°C. For RNA-seq, *U. maydis*-, *S. reilianum-* and rUSH-infected Golden Bantam maize seedlings were used. For *S. reilianum* infection assays maize seedlings of the cultivar Gaspe Flint were infected. For infection, a 1:1 mixture of compatible mating partners or the solopathogenic strain SG200 was used (OD_600_ of 1). For rUSH infections and microscopy, 0.1% Tween was added prior to infection. The disease symptoms for *U. maydis* were evaluated 12 dpi and for *S. reilianum* 8 wpi.

### Staining and microscopy

To observe the formation of appressoria-like structures, 20-22 hpi infected maize leaves were incubated in calcofluor white staining solution (100 µg ml−1; in 0.2 M Tris-HCl; pH 8.0) for 1 min and briefly rinsed with water before observation using the DAPI filter on the Nikon Eclipse Ti Inverted Microscope (Lanver *et al*., 2014). To visualize fungal growth in infected leaves, WGA-AF488 (Wheat Germ Agglutinin, Alexa Fluor 488) and propidium iodide were used to stain fungal and plant cell walls, respectively, as previously described by Doehlemann *et al*. (2009). For microscopy a Nikon Eclipse Ti Inverted Microscope was utilized alongside the Nikon NIS-ELEMENTS software (Düsseldorf, Germany). Images were captured using a HAMAMATSU camera. To visualize fluorescent proteins, the Leica TCS SP8 Confocal Laser Scanning Microscope (Leica, Bensheim, Germany) was employed. GFP was excited at 488 nm and detected within the 490-540 nm range, while mCherry was excited at 561 nm and detected from 580 to 660 nm. The microscopy images were processed using the Leica LAS X.Ink software.

### DNA and RNA preparation and qRT-PCR

Three maize infections were conducted from three independent fungal cultures. 4 cm-long sections (1 cm below the infection side) of the third leaf were collected from at least 12 individual plants. At 20 hpi, ∼2 cm-long leaf sections were used and liquid latex was applied to the infected maize leaves, dried, and peeled off for RNA extraction. At the 3 dpi and 6 dpi the plant material was used as described above. The frozen plant tissue and latex were ground into a fine powder using liquid nitrogen. Total RNA was extracted by using TRIzol (Thermo Fisher, Waltham, USA) according to the manufacturer’s protocol. Subsequently, a DNase I digest was performed (Thermo Fisher) and the samples were sent to Novogene (UK) for RNA-seq. For qRT-PCR, cDNA was synthesized using RevertAid First Strand cDNA Synthesis kit (Thermo Fisher). The qRT-PCR was performed using a GoTaq qPCR mix (Promega) and a CFX96 Real-Time PCR Detection System (BioRad). DNA extraction of *U. maydis* and *S. reilianum* cultures was prepared using smut lysis buffer (10 mM tris HCl (pH 8.0), 100 mM NaCl, 1 mM Na2-EDTA, 1% SDS, 2% Triton). For biomass quantification, the DNA was isolated using maize extraction buffer (0.1 M Tris-HCl, 0.05 M EDTA, 0.5 M NaCl, 1.5% SDS; after autoclaving addition of 0.3% β-mercaptoethanol) and subsequently purified using a MasterPure Complete DNA and RNA Purification Kit (Epicenter, Madison, USA). For biomass quantification 150 ng of DNA was used. To determine the ratio between fungal *peptidylprolyl isomerase* (*ppi*) and maize *GAPDH* for quantification: 2^ΔCt^ (Ct*^ZmGAPDH^* - Ct*^Umppi^*) and for relative gene expression 2^ΔCt^ (Ct*^Umppi^* – Ct*^GOI^*) were calculated. For statistical analysis, a student’s t-test was conducted.

### RNA-sequencing data analysis

The RNA-seq was conducted by Novogene. RNA libraries were prepared using an Illumina TruSeq paired-end sequencing (Illumina, SanDiego, CA, USA) performed on a HiSeq4000 platform. Reads of three biological replicates were filtered using the Trimmomatic software 0.39 and standard settings and mapped to a reference assembly using BOWTIE2 (version 2.3.4.1) (Langmead & Salzberg, 2012). The reference genomes of *U. maydis* (Kämper et al., 2006) and *S. reilianum* (Schirawski *et al*., 2010) were combined before mapping to eliminate a mapping bias to the wrong species. Reads were counted to *U. maydis* and *S. reilianum* using HTseq-count (version 2.0.4) (Anders *et al*., 2015). The edgeR package was used for statistical analysis of differential gene expression (counts per million, CPM) and Excel was used to calculate the transcripts per million (TPM) normalized to the different gene lengths between the species. Afterward, one-to-one orthologs were determined (Zuo et al. 2021). For the comparison between SRZ1xSRZ2 (SR) and FB1xFB2 (UM), the fold change FC(SR/UM) was calculated by dividing the *S. reilianum* transcripts per million (TPM) by *U. maydis* TPM of the wild types. The same analysis was conducted for the orthologous genes in rUSH (Differentially expressed genes: log_2_FC>1 or <−1, p<0.05). Differentially expressed effector genes (log_2_FC>1 or <-1; p<0.05) were grouped using KMeans clustering with one minus Pearson’s correlation. The total expression of each ortholog at 3 and 6 dpi was compared between WT and rUSH, leading to the creation of 8 clusters with the following patterns at 6 dpi: C1: higher in SR in WT, higher UM in rUSH (ortholog-specific), C2: no difference in WT, higher in SR in rUSH (ortholog-specific), C3: higher in UM in WT, downregulated in rUSH (*trans*-regulated), C4: higher in SR in WT and rUSH (*cis*-regulated), C5: higher in UM in WT and rUSH (*cis*-regulated), C6: higher in SR in WT, not differentially expressed in rUSH (*trans*-regulated), C7: higher in UM in WT, higher in SR in rUSH (reverse expression), C8: high in SR in WT, higher in SR in rUSH (ortholog-specific expression). ShinyGO v0.66: Gene Ontology Enrichment Analysis + more was used for the GO term enrichment analysis of maize genes upregulated in rUSH_OE_*Umhdp2* vs. rUSH (http://bioinformatics.sdstate.edu/go65/).

## Acknowledgements

This project has received funding from the European Research Council (ERC) under the European Union’s Horizon 2020 research and innovation program (grant agreement No. 771035) as well as from the Cluster of Excellence on Plant Sciences (CEPLAS) funded under Germany’s Excellence Strategy—EXC 2048/1—project ID: 390686111.

## Author contributions

GD and WZ supervised the project. GD, WZ and JW designed the experiments. JW and WZ conducted the experiments, TW generated FB1_*Sra1b1*. JW wrote the manuscript with contributions from all authors.

## Data availability

Source data are provided in this paper. All data supporting the findings of this study that are not directly available within the paper (and its supplementary data) will be upon reasonable request available from the corresponding authors (GD, JW). Raw data of RNA-seq analysis are publicly accessible in the NCBI Gene Expression Omnibus (accession number: XXX).

## Supporting information

**Figure S1:** Generation of SRZ1_*Uma1b1*

**Figure S2:** Downregulation of *U. maydis* effector genes is not caused by RNAi machinery of *S. reilianum*

**Figure S3:** Differentially expressed one-to-one orthologs in rUSH compared to FB1xSRZ2.

**Figure S4:** CRISPR/Cas9a frameshift mutants of putative transcription factors revealed no effect on *U. maydis’* virulence

**Supplementary Table S1:**

- RNA-seq (WT/rUSH)
- List of U. maydis effector candidates
- Downregulated *U. maydis* effector candidates (5-fold cut-off)
- Upregulates *S. reilianum* effector candidates (2-fold cuf-off)
- Transcription factor candidates
- Regulation of effector candidates (WT/rUSH)
- RNA-seq rUSH vs. FB1xSRZ2 (UM)
- RNA-seq rUSH vs. FB1xSRZ2 (SR)
- RNA-seq rUSH_OE_*Umhdp2* vs. rUSH (UM)
- RNA-seq rUSH_OE_*Umhdp2* vs. rUSH (ZM)
- Enrichment ZM up - rUSH_OE_*Umhdp2*

