## Supplemental Figures for "A recombinant hybrid provides insights into gene regulation, pathogenesis and tumorigenesis of phytopathogenic smut fungi"

### Supplementary figures

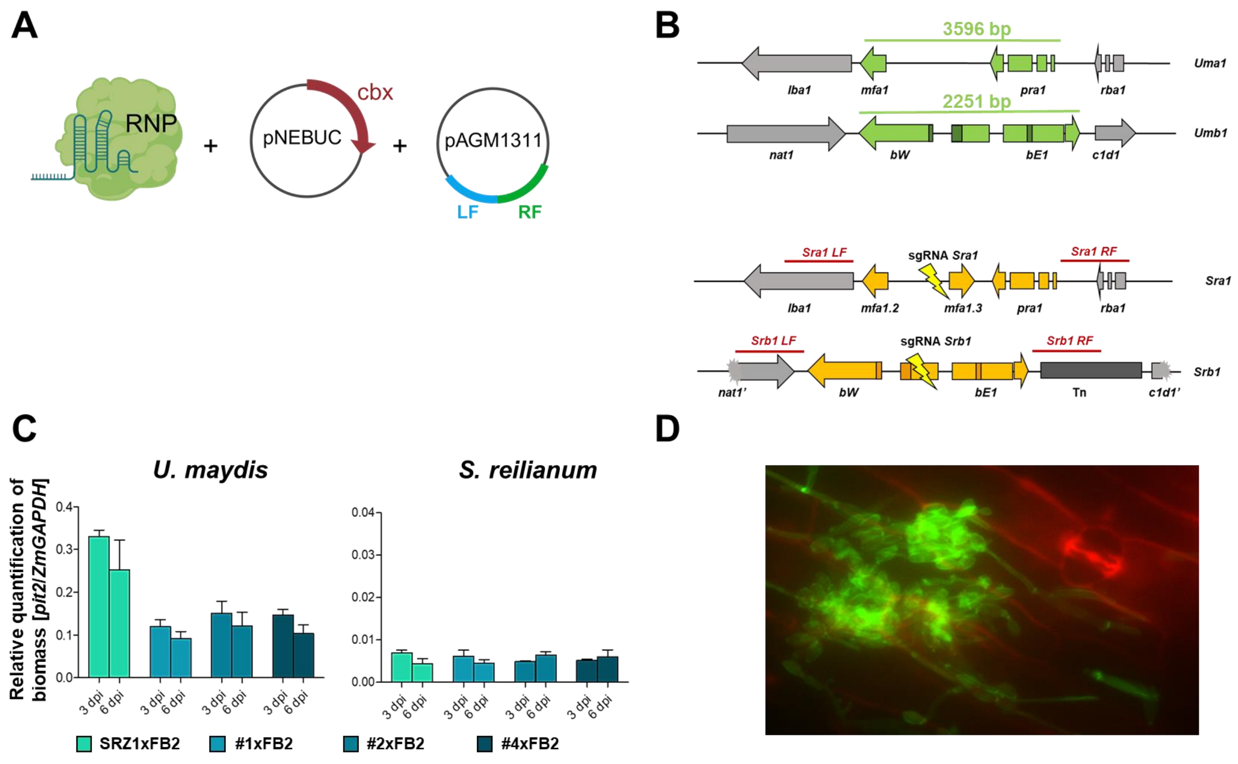


**Figure S1: Generation of SRZ1_*Uma1b1*. (A)** SRZ1_*Uma1b1* was generated using ribonucleoprotein (RNP)-mediated transformation with a donor template (Werner *et al*., 2024) and an auto-replicating plasmid (pNEBUC). **(B)** Schematic overview of the *a1* and *b1* locus of *S. reilianum* (orange) and *U. maydis* (green). sgRNAs are indicated in yellow. 1 kb flanks used for the donor template are marked in red. **(C)** qRT-PCR of SRZ1xFB2, SRZ1_*Uma1b1*#1xFB2 (#1xFB2), SRZ1_*Uma1b1*#2xFB2 (#2xFB2) and SRZ1_*Uma1b1*#4xFB2 (#4xFB2) for quantification of fungal biomass using *Umppi* or *Srppi*, respectively, and *GAPDH* of maize. **(D)** WGA staining of #1xFB2 at 6 dpi revealed a clump-like structure on the plant surface.

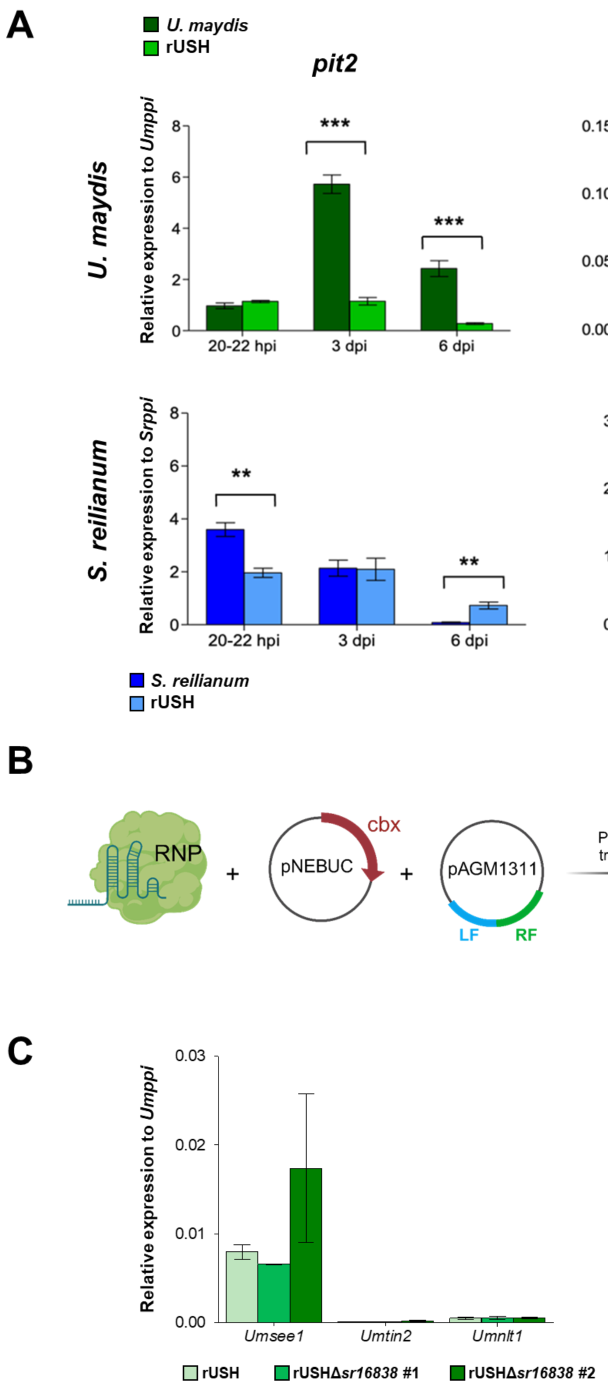


**Figure S2: Downregulation of *U. maydis* effector genes is not caused by RNAi machinery of *S. reilianum*. (A)** Gene expression of the one-to-one effector orthologs *pit2* and *see1* in rUSH. Gene expression of *pit2* and *see1* were measured and normalized to the *ppi* gene of *U. maydis* or *S. reilianum*, respectively. Significant differences were calculated based on students t-test (* = p< 0.05, ** = p< 0.01, *** =p<0.001). **(B)** Schematic illustration of RNP-mediated *S. reilianum* transformation. *sr16838* was deleted in the SRZ2 to test, whether the dicer gene is responsible for the downregulation of *U. maydis* effector orthologs in rUSH. Therefore, 1 kb upstream and 1 kb downstream of the cds were amplified and using Gibson assembly cloned into pAGM1311 linearized with XbaI and EcoRI. The resulting donor template was used together with a sgRNA cutting approximately in the middle of the cds of *sr16838* in an

RNP and the autoreplicating plasmid pNEBUC. Cbx was used for selection. **(C)** Relative Expression of *Umnlt1*, *Umsee1* and *Umtin2* in rUSHΔ*sr16838*. Maize leaves were infected with rUSH, rUSHΔ*sr16838*#1, and rUSHΔ*sr16838*#2 with a final OD of 1 and addition of 0.1% Tween. After 3 dpi 4-cm leaf sections were harvested and further processed. Using qRT-PCR, gene expression was measured relative to *Umppi* (green). Error bars (standard deviation) were calculated from four biological replicates.

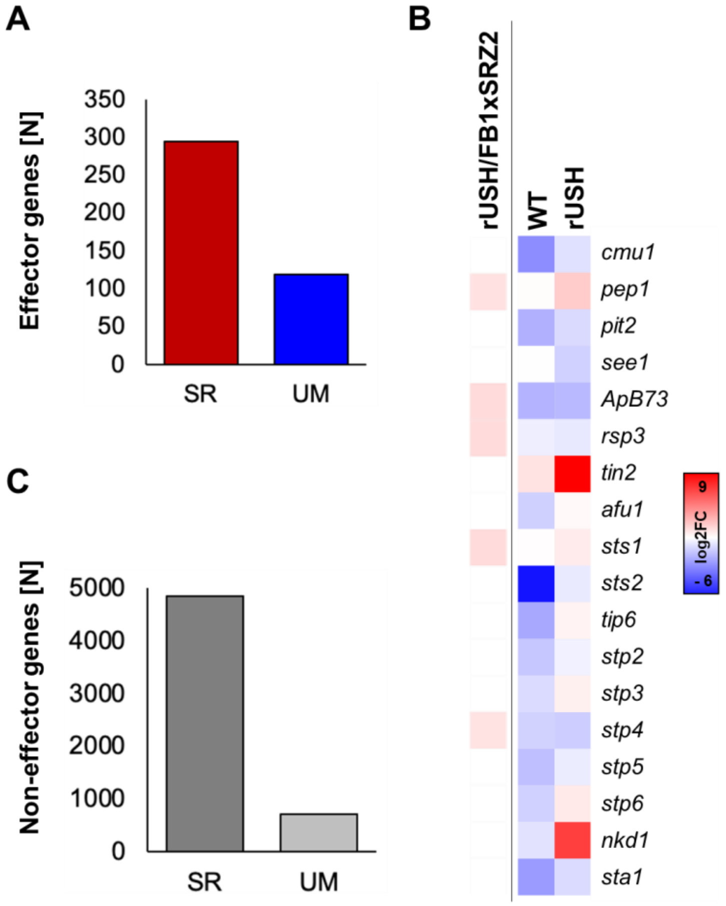


**Figure S3: Differentially expressed one-to-one orthologs in rUSH compared to FB1xSRZ2.** RNA-seq analysis was conducted and differentially expressed one-to-one orthologs ($log_2FC>1$, $p<0.05$; Zuo *et al*., 2021) were grouped into effector genes **(A)** and non-effector genes **(C)** and compared between *U. maydis* (UM) and *S. reilianum* (SR) in rUSH and FB1xSRZ2. **(B)** Differentially expressed effector genes in rUSH vs. FB1xSRZ2; red indicates an upregulation of the *U. maydis* ortholog in rUSH. For WT (WT_SR/WT_UM) and rUSH (rUSH_SR/rUSH_UM) the SR TPM were divided by UM TPM (blue: *U. maydis* ortholog > *S. reilianum* ortholog, red: *S. reilianum* ortholog > *U. maydis* ortholog).

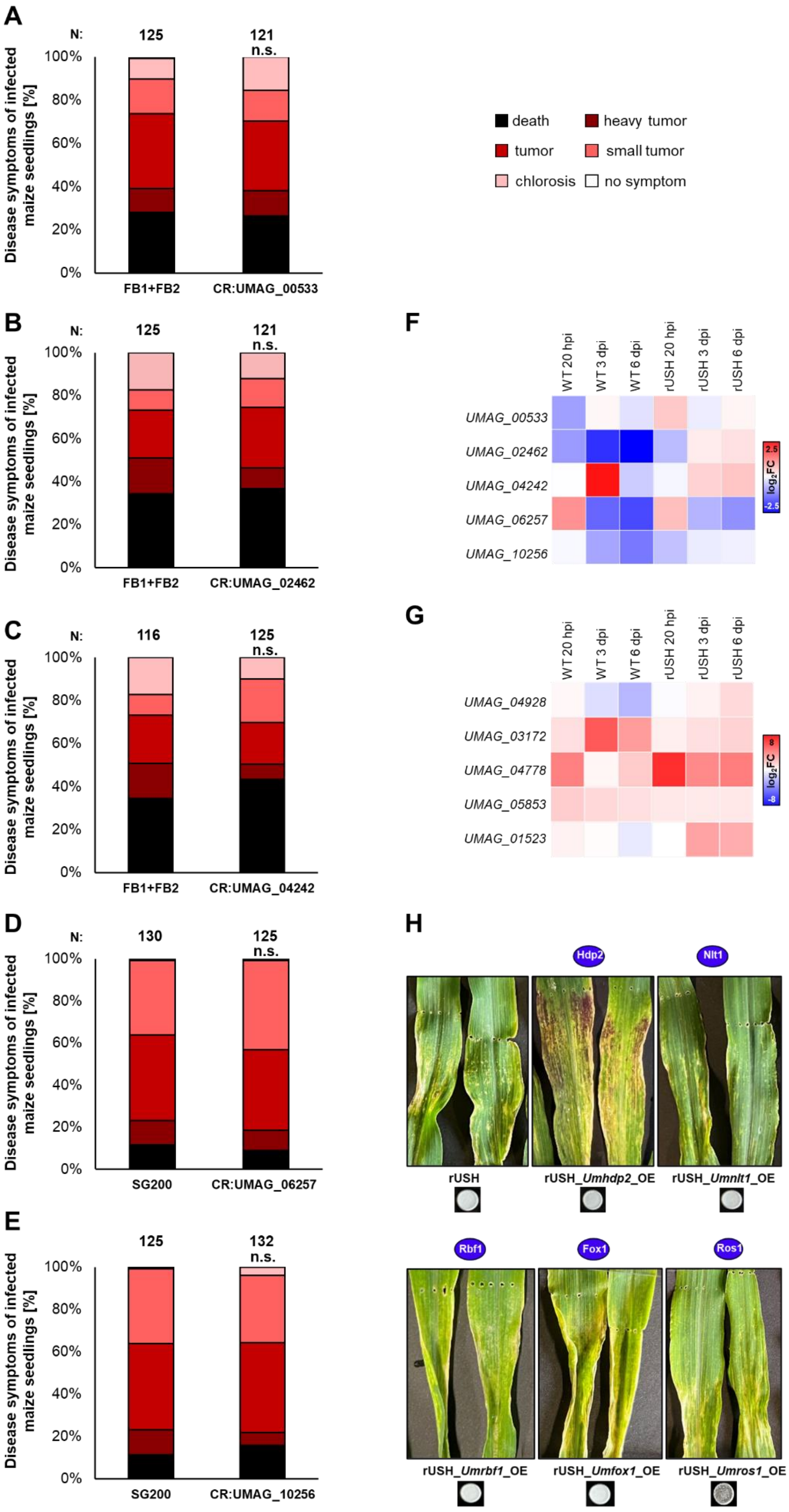
A
N:
125
121
n.s.
Disease symptoms of infected maize seedlings [%]
100%
80%
60%
40%
20%
0%
FB1+FB2
CR:UMAG_00533
death
heavy tumor
tumor
small tumor
chlorosis
no symptom
B
125
121
FB1+FB2
CR:UMAG_02462
C
116
125
FB1+FB2
CR:UMAG_04242
D
130
125
SG200
CR:UMAG_06257
E
125
132
SG200
CR:UMAG_10256
F
WT 20 hpi
WT 3 dpi
WT 6 dpi
rUSH 20 hpi
rUSH 3 dpi
rUSH 6 dpi
UMAG_00533
UMAG_02462
UMAG_04242
UMAG_06257
UMAG_10256
log2FC
2.5
-2.5
G
UMAG_04928
UMAG_03172
UMAG_04778
UMAG_05853
UMAG_01523
8
-8
H
Hdp2
Nlt1
rUSH
rUSH_Umhdp2_OE
rUSH_Umnlt1_OE
Rbf1
Fox1
Ros1
rUSH_Umrbf1_OE
rUSH_Umfox1_OE
rUSH_Umros1_OE

**Figure S4: CRISPR/Cas9 frameshift mutants of putative transcription factors revealed no effect on *U. maydis*' virulence.** Infection assays of 7-days old maize seedlings (cultivar: Golden Bantam) infected with CR-mutants in FB1xFB2 background: **(A)** *UMAG_00533*, **(B)** *UMAG_02462* and **(C)** *UMAG_04242* or in SG200 background: **(D)** *UMAG_06257*, **(E)** *UMAG_10256*. **(F)** Heatmap of differentially expressed putative transcription factors (*UMAG_00533*. *UMAG_02462*, *UMAG_04242*, *UMAG_06257*, *UMAG_10256*). Disease symptoms were scored 12 dpi. **(G)** Heatmap of *Umhdp2* (*UMAG_04928*), *Umrbf1* (*UMAG_03172*), *Umnlt1* (*UMAG_04778*), *Umros1* (*UMAG_05853*) and *Umfox1* (*UMAG_01523*). **(F+G)** $Log_2FC$ was calculated by division of *S. reilianum* TPM by *U. maydis* TPM for WT (wild type) and rUSH at 20-22 hpi, 3 dpi and 6 dpi. **(H)** Infection assay of TF overexpressions in rUSH. *Umhdp2*, *Umnlt1*, *Umrbf1*, *Umfox1*, and *Umros1* were overexpressed in FB1_*Sra1b1* and infected together with SRZ2 (mixed 1:1, final OD of 1) in 7 days old maize seedlings. Pictures were taken at 6 dpi.
